# A contraction stress model of hypertrophic cardiomyopathy due to thick filament sarcomere mutations

**DOI:** 10.1101/408294

**Authors:** Rachel Cohn, Ketan Thakar, Andre Lowe, Feria Ladha, Anthony M. Pettinato, Emily Meredith, Yu-Sheng Chen, Katherine Atamanuk, Bryan D. Huey, J. Travis Hinson

## Abstract

Thick filament sarcomere mutations are the most common cause of hypertrophic cardiomyopathy (HCM), a disorder of heart muscle thickening associated with sudden cardiac death and heart failure, with unclear mechanisms. We engineered an isogenic panel of four human HCM induced pluripotent stem cell (iPSc) models using CRISPR/Cas9, and studied iPSc-derived cardiomyocytes (iCMs) in 3-dimensional cardiac microtissue (CMT) assays that resemble *in vivo* cardiac architecture and biomechanics. HCM mutations result in hypercontractility in association with prolonged relaxation kinetics in proportion to mutation pathogenicity but not calcium dysregulation. RNA sequencing and protein expression studies identified that HCM mutations result in p53 activation secondary to increased oxidative stress, which results in increased cytotoxicity that can be reversed by p53 genetic ablation. Our findings implicate hypercontractility as an early consequence of thick filament mutations, and the p53 pathway as a molecular marker and candidate therapeutic target for thick filament HCM.

## Introduction

Hypertrophic cardiomyopathy (HCM) is a human disorder that affects 1 in 500 individuals with uncertain mechanisms^1^. Patients with HCM are diagnosed by the presence of unexplained left ventricular hypertrophy (LVH) with preserved systolic contractile function^2^. In young athletes, HCM manifests as a common cause of sudden cardiac death; while in adults, HCM is associated with heart failure that may progress to require cardiac transplantation^3^. Over the last few decades, the genetic basis of HCM has been demonstrated by inheritance of autosomal dominant mutations in components of the force-producing sarcomere^4^. About two-thirds of HCM patients harbor heterozygous mutations in one of two sarcomere genes: β-myosin heavy chain (MHC-β is encoded by *MYH7*) or myosin-binding protein C (MYBPC3)^4^. Along with titin, MHC-β and MYBPC3 are located in the thick filament where ATP hydrolysis by MHC-β is coupled to force generation through interactions with the actin-rich thin filament (fig.1A). A prevailing model suggests that HCM mutations alter cardiac force generation through dysregulation of calcium handling^5, 6^. Whether MYBPC3 and MYH7 mutations result in HCM by shared or heterogeneous mechanisms also remains unclear.

**Figure 1.**
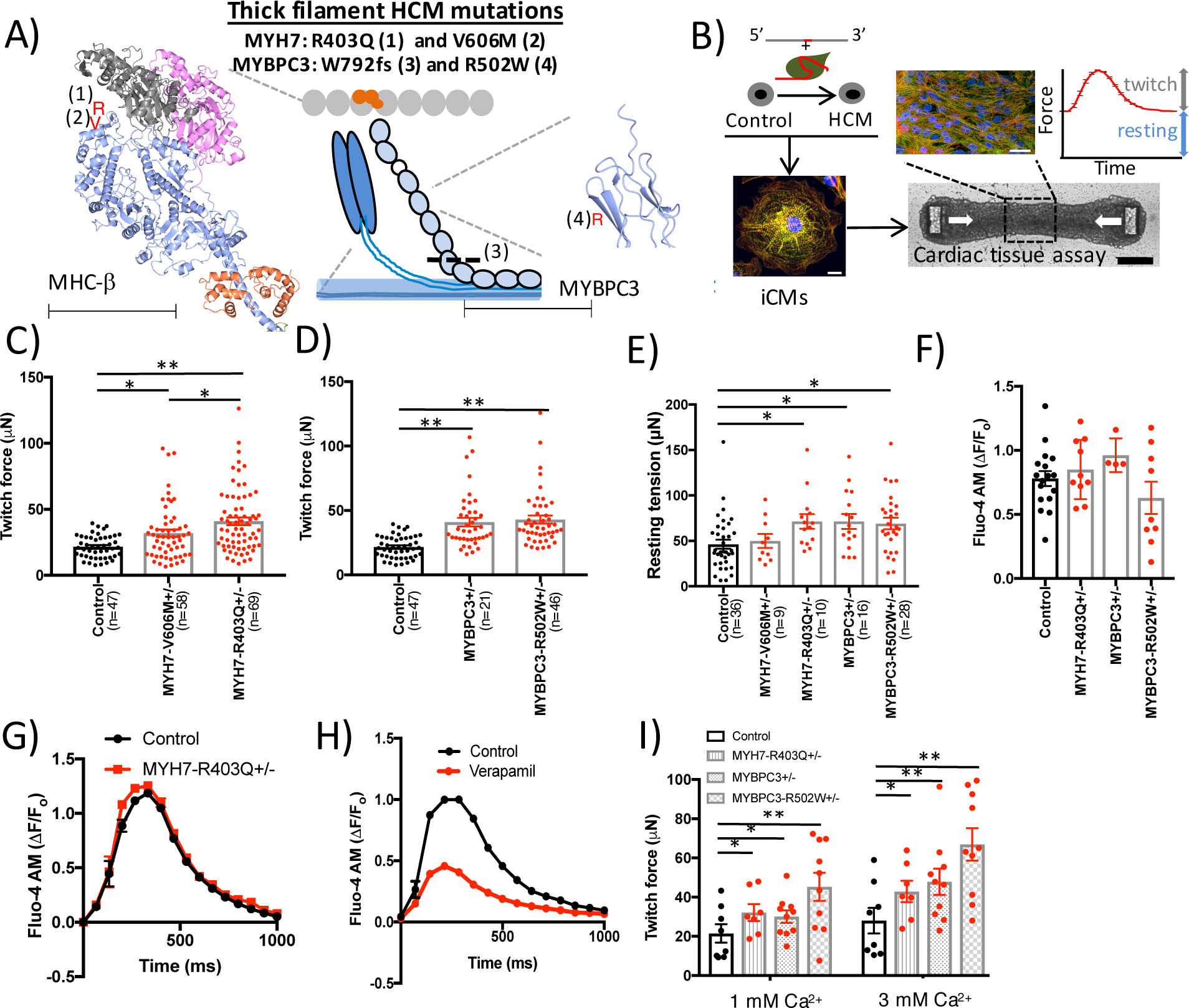
Human iPSc-derived CMT models with thick filament HCM mutations result in hypercontractility. (**A**) A representation of the sarcomere is shown that includes thick filament components β-myosin heavy chain (MHC-β; blue globular heads connected to thin rods) and myosin-binding protein C (MYBPC3; chain of light blue ovals); and thin filament components actin (gray ovals) and the troponin complex (orange ovals). Location of mutations are decorated on the crystal structures of MHC-β-S1 (blue ribbon; left) and a domain of MYBPC3 (blue ribbon; right)^47^. Note: MHC-β-S1 is shown interacting with two actin molecules (gray and pink ribbons) and a regulatory light chain (orange ribbon). For MYH7, R403Q is identified by a red R (1), and V606M is denoted by a red V (2). For MYBPC3, the location of the truncation W792fs is denoted by a dashed line (3), and R502W is denoted by a red R (4). Scale bar=62.5Å (MHC-β;) and 31Å (MYBPC3). (**B**) Experimental outline of isogenic HCM model generation using gRNA/Cas9 complex and ssODN to introduce HCM mutations into a control iPSc line. IPScs are then differentiated to produce iCMs (scale bar=10μm) that are combined with fibroblasts and an extracellular matrix slurry for CMT production (white arrows depict direction of contraction; top panel-scale bar=25μm and bottom panel-scale bar=200μm)). Both tissue twitch force and resting tension are quantified as well as CMT sarcomere structure by immunofluorescence. (**C**) Maximum twitch force from CMTs generated from control, MYH7-V606M^+/-^ and MYH7-R403Q^+/-^ iCMs. (**D**) Maximum twitch force from CMTs composed of control, MYBPC3^+/-^ and MYBPC3-R502W^+/-^ iCMs. (**E**) Resting tension produced by HCM CMTs compared to controls. (**F**) Quantification of calcium transients (ΔF/F_o_) measured in HCM and control CMTs stained with Fluo-4 while pacing at 1Hz (see representative tracing **(G)**). **(H)** Representative calcium transient tracing of control iCMs treated with verapamil or carrier control. **(I)** Dependence of maximum twitch force generated by HCM and control CMTs from extracellular calcium concentration. Significance assessed by ANOVA (C-F, I; *all p<0.05 and **all p<0.001); data are means +/- SEM (error bars) (C-I).

Recent functional studies of thick filament HCM mutations in reconstituted sarcomere and cardiomyocyte assays have supported both gain- and loss-of-force production models of HCM pathogenesis, thus suggesting that changes in force production may not be a shared consequence of HCM mutations. For example, MYH7-R453C (arginine 453 substituted with cysteine) increased while MYH7-R403Q (arginine 403 substituted with glutamine) decreased force production in reconstituted actomyosin motility assays^7, 8^. Equally puzzling, contractile studies of single cardiomyocytes from MYH6-R403Q^+/-^ mouse models, which recapitulate LVH and fibrosis *in vivo*^9^, have produced similarly conflicting results for the identical mouse model and strain^10, 11^. Human patient-specific induced pluripotent stem cell (iPSc) HCM models of MYH7-R663H (arginine 663 substituted with histidine) have recapitulated some features of HCM including cellular enlargement and altered calcium handling^6^, but mechanical phenotypes of HCM iPSc models have not been comprehensively studied.

The apparent difficulty in establishing the pathogenesis of HCM has been attributed in part to: 1) multiprotein assembly limitations that hinder sarcomere analysis, 2) mouse models that express distinct sarcomere components compared to humans (e.g. *MYH6* instead of *MYH7*), 3) the lack of isogenic iPSc-derived cardiomyocyte cell lines to control for genetic and epigenetic variation and 4) the absence of biomimetic 3-dimensional (3d) human cardiac tissue functional assays. Here, we set out to address these limitations by combining genetic engineering tools to generate a series of scarless *MYH7* and *MYBPC3* HCM mutations in human isogenic iPScs that are differentiated to cardiomyocytes (iCMs) that express human sarcomere contractile components. We generated 3d cardiac microtissues (CMTs; fig.1B) to identify mechanical consequences of HCM mutations in combination with molecular assays to interrogate insights into HCM pathogenesis.

## Results

### Generation of HCM iPSc and CMT models using CRISPR/Cas9

We began by identifying two HCM mutations in *MYH7*, R403Q and V606M (valine 606 substituted with methionine), which cause autosomal dominant HCM in both humans^9, 12^ and mice^13, 14^. Both mutations are located in subfragment 1 (S1) of MHC-β near the actin-interacting domain (fig.1A), but R403Q leads to a more severe cardiomyopathy compared to V606M^14^. Because MHC-β physically interacts with MYBPC3^4^, we hypothesized that these mutations could lead to HCM by a shared mechanism.

We next selected a common pathogenic truncation mutation in *MYBPC3* (clinvar.com), a guanine insertion that leads to tryptophan 792 substituted with valine followed by a frameshift (Trp792ValfsX41; MYBPC3^+/-^), and a common pathogenic missense mutation, arginine 502 substituted with tryptophan (R502W) that is found in up to 2.4% of HCM patients (fig.1A)^15^. We generated scarless, isogenic iPSc models of these four mutations with CRISPR (clustered regularly interspaced short palindromic repeats) technology using delivery of a single optimized guide RNA, Cas9 nuclease and a single-stranded oligodeoxynucleotide (ssODN) repair template (fig.1B, table S1). We chose an isogenic approach to focus our investigation on the direct functional consequences of the four thick filament mutations, while controlling for background epigenetic and genetic variation. To generate the four iPSc models, we screened 1,111 clones by Sanger sequencing (table S1). After screening for off-target genome editing loci by sequencing (crispr.mit.edu) and karyotype abnormalities by virtual karyotyping using arrays, we directly differentiated iPScs^16^ followed by metabolic enrichment^17^ to generate purified iCMs. Finally, we combined iCMs with tissue-forming fibroblasts and an extracellular matrix slurry to generate a 3d CMT assay that recapitulates native cardiac architecture and mechanics, which has been adapted from prior assays applied to study contractility phenotypes of dilated and PRKAG2 cardiomyopathy iPSc models^18-20^

### Four thick filament HCM mutations result in hypercontractility in CMT assays

To study the mechanical consequences of *MYH7* and *MYBPC3* mutations in a biomimetic context, we optimized our previously studied CMT assay^18, 20^ to increase sarcomere gene expression of thick filament transcripts. In particular, *MYH7:MYH6* is upregulated in the developing human heart^21^ and iCMs express fetal to neonatal stage transcript levels. By increasing the size of the microfabricated tissue gauges (μTUG) including a proportional increase in cantilever dimensions and spring constant (fig.S1A), we improved CMT durability from four days to ten days (fig.S1B), which was associated with a 42% increase in *MYH7*:*MYH6* expression (fig.S1C, left panel) and 630% increase in cardiac troponin I (*TNNI3*) expression (fig.S1C, right panel). For all CMT studies, we measured contractility parameters on day seven after tissue compaction has completed and force production has plateaued (fig.S1D).

CMTs generated from iCMs with MYH7-R403Q^+/-^ (fig.S1E), generated 40.9 μN twitch force compared to 21.7 μN in isogenic controls (fig.1C). We then tested whether the increased twitch force generated by R403Q^+/-^ was secondary to changes in sarcomere isoform expression or clonal variation. *TNNI3* expression^22^, a transcript marker that is related to cardiomyocyte maturation, was unchanged (fig.S2A, B)^22^. Increased contraction force was also similar between two independent R403Q^+/-^ clones (fig.S2C). Moreover, CMT cross-sectional area did not differ between HCM mutations and isogenic controls (fig.S2D). We next tested whether CMT assays could predict pathogenicity of *MYH7* mutations by testing CMTs generated from iCMs with the less pathogenic *MYH7* variant V606M^+/-^, which is also located in the actin-binding domain of S1 (fig.S1F). V606M^+/-^ CMTs generated 31.9 μN twitch force compared to 21.7 μN in isogenic controls (fig.1C), which was less than R403Q^+/-^. We conclude that *MYH7* variant pathogenicity positively correlates with maximum contraction force in CMT assays. We also generated two *MYBPC3* mutant CMT models to compare to *MYH7* models. MYBPC3^+/-^ and MYBPC3-R502W^+/-^ CMTs generated 41.0 μN and 42.9 μN twitch force compared to 21.7 μN in isogenic controls, respectively (fig.1D). Because sarcomere function also contributes to twitch-independent contraction force, or resting tension, we measured this parameter in CMTs. In parallel to twitch force changes, resting tension was increased in all HCM models except for MYH7-V606M^+/-^ (fig.1E). Because V606M^+/-^ results in a mild phenotype, we did not further characterize this variant.

To characterize the basis of hypercontractility, we measured calcium handling in CMTs by optical imaging and calcium-dependent fluorescent dyes. Surprisingly, HCM-associated hypercontractility did not appear to be related to changes in calcium delivery to the myofilament as calcium transients were unaffected in all HCM mutations tested (fig.1F, G). This is distinct from CMT treatment with verapamil, a voltage-gated calcium channel blocker, which resulted in diminished CMT calcium transients as expected (fig.1H). Because the increase in resting tension could be secondary to myofilament activation in the setting of increased resting calcium levels, we stained iCMs with Indo-1 and measured both resting and caffeine-induced activation. Indo-1 signal was unchanged (fig.S2E, F). Finally, we tested CMT myofilament calcium sensitivity by varying the extracellular calcium concentration from 1 mM to 3 mM and quantified CMT function. We found that at both low and high calcium, HCM CMTs resulted in hypercontractility (fig.1I). These data support a model whereby thick filament HCM mutations result in a state of hypercontractility that is independent of changes in calcium handling.

We also measured HCM-associated contraction and relaxation kinetics by quantifying maximum contraction velocity, contraction time and relaxation half-time across all HCM mutations. Compared to controls, HCM CMTs exhibited a 79-121% increased maximum contraction velocity (fig.2A, B), but without changes in contraction time (fig.2C). Relaxation half-time (t_1/2_) was prolonged (fig.2D), which further supports that impaired relaxation is a consequence of HCM thick filament mutations, which has also been documented in HCM patients^23^. These kinetic changes are distinct from kinetic changes induced by omecamtiv mecarbil^24^, a direct myosin activator that prolongs contraction time.

**Figure 2.**
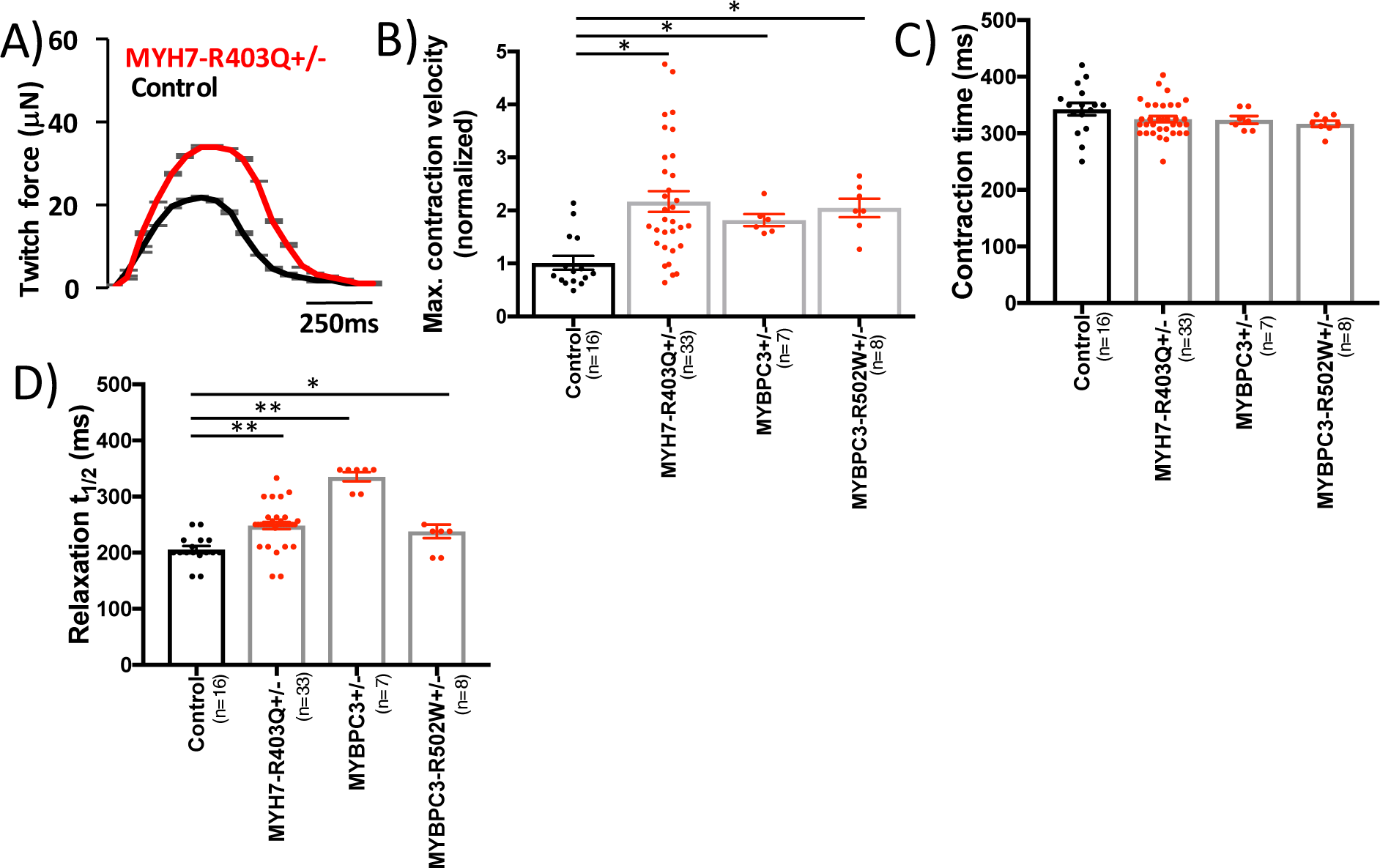
Tissue contraction and relaxation kinetics are altered by HCM mutations. (**A**) Representative twitch force tracings from MYH7-R403Q^+/-^ CMT models compared to isogenic controls. (**B**) Normalized maximum contraction velocity, **(C)** contraction time and **(D)** relaxation halftime (t_1/2_) for MYBPC3^+/-^, MYBPC3-R502W^+/-^ and MYH7-R403Q^+/-^ compared to controls. Significance was assessed by ANOVA (B-D; *all p<0.05 and **all p<0.001); data are means +/- SEM (error bars) (A-D).

### Reduction of hypercontractility by calcium channel blockers and direct myosin inhibitors

We generated CMTs from the severe HCM mutation MYH7-R403Q^+/-^, and assessed changes in twitch force and resting tension after treatment with two candidate small molecules. We started by evaluating one of the most commonly prescribed therapies for HCM patients, the voltage-dependent calcium channel blocker verapamil^25, 26^. In accord with changes in calcium transients (fig.1H), the addition of verapamil reduced twitch tension by 31.7% in R403Q^+/-^ CMTs (fig.3A). We also tested the direct myosin inhibitor blebbistatin to assess whether myosin inhibitors may similarly reduce HCM-associated hypercontractility in R403Q^+/-^ CMTs. Since blebbistatin binds to MHC-β distinct from residues near R403Q, and does not interfere with actin-binding or actomyosin disassociation^27^, we hypothesized that this molecule may also reduce twitch force. Similar to verapamil, blebbistatin similarly reduced twitch force by 35.3% (fig.3B). Because blebbistatin inhibits myosin by a calcium-independent mechanism and resting calcium levels were not altered by HCM mutations, we hypothesized that only blebbistatin could normalize both twitch and resting tension induced by HCM mutations. Indeed, blebbistatin but not verapamil reduced resting tension by 10.2% (fig.3C). These data obtained from R403Q^+/-^ CMTs suggest that therapeutic agents that reduce both twitch and resting forces may be more efficacious for HCM patients, which may explain why verapamil has been observed to have limited clinical efficacy in some HCM patients^28^.

**Figure 3.**
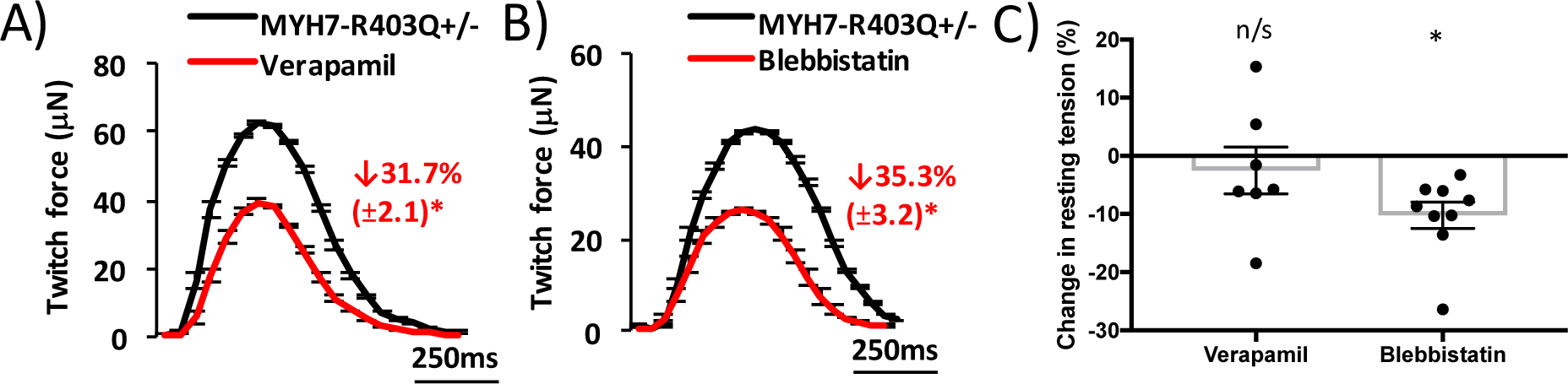
Reduction of hypercontractility in MYH7-R403Q^+/-^ tissues by small molecules. Representative tracings and % change in twitch force for MYH7-R403Q^+/-^ CMTs treated with carrier control (black tracing) compared to **(A)** verapamil (0.5μM; red tracing) or **(B)** blebbistatin (10μM; red tracing). **(C)** Comparison of the effects of verapamil and blebbistatin on resting tension in MYH7-R403Q^+/-^ CMTs. Significance was assessed by Student’s t-test (A-C; *all p<0.05); data are means +/- SEM (error bars) (A-C).

### MYH7-R403Q^+/-^ CMTs exhibit myofibrillar disarray and iCM hypertrophy

To determine whether HCM mutations affect sarcomere structure in CMTs, we fixed CMTs and stained sarcomeres using antibodies to Z-disc protein alpha actinin with DAPI co-stain (fig.4A). We again focused on R403Q^+/-^ because this mutation is highly pathogenic in humans and in our mechanical assays. Z-disk analysis of R403Q^+/-^ tissues demonstrated increased Z-disk angular dispersion (figs.4B, S3A)), which is a measure of Z-disk disarray. This result is consistent with myofibrillar disarray observed in HCM patients^29^. Sarcomere length and cell number were not different (fig.S3B, C) in R403Q^+/-^ CMTs compared to isogenic controls. To assess iCM size, we fixed and immunostained R403Q^+/-^ iCMs (fig.4C) because the analysis of individual iCMs is infeasible within our CMT assays. Single R403Q^+/-^ iCMs demonstrated increased iCM cell area (fig.4D). We next measured levels of candidate hypertrophy signaling pathways including mitogen-activated protein kinase pathways, AKT and CAMKII as these have been implicated in cardiomyocyte hypertrophy *in vivo* and in iCMs^18, 30^. In proportion to increased iCM cell area, lysates from R403Q^+/-^ iCMs had elevated levels of phosphorylated ERK2 and AKT (fig.4E-G), but not p38, JNK or CAMKII (fig.S3D). In summary, R403Q^+/-^ mutations result in sarcomere disorganization in CMT assays, and iCM hypertrophy in parallel with increased ERK2 and AKT signaling.

**Figure 4.**
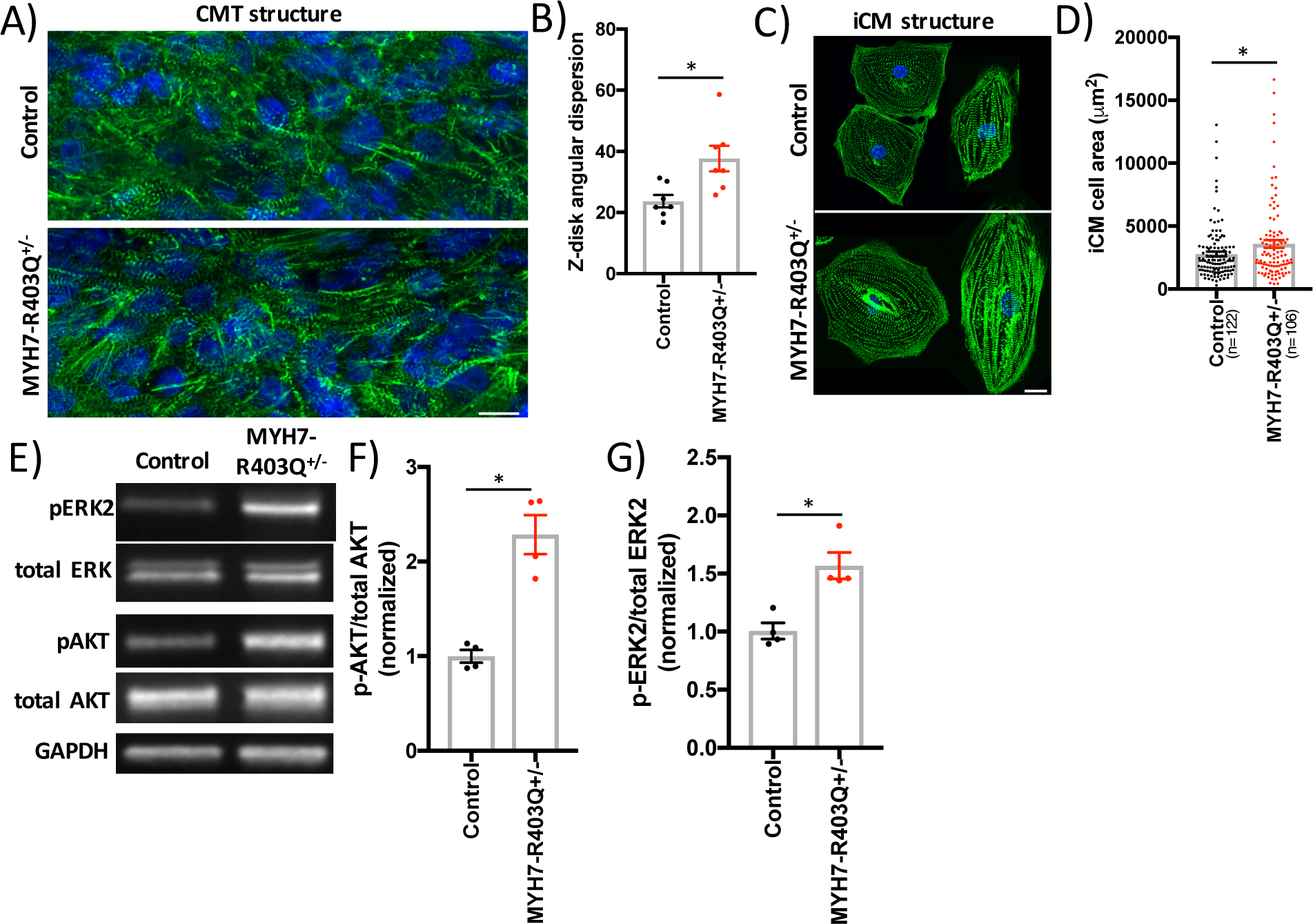
CMT and iCM structural and molecular signaling changes in MYH7-R403Q^+/-^ models. (**A**) Representative confocal image of CMTs generated from MYH7-R403Q^+/-^ and control iCMs and decorated with antibodies to cardiac alpha actinin (green) and co-stained with DAPI (blue; scale bar=10μm). **(B)** Quantification of sarcomeric Z-disk angular dispersion obtained from analysis of confocal regions of interest from CMTs decorated with antibodies to alpha actinin. **(C)** Representative confocal images of single MYH7-R403Q^+/-^ and control iCMs decorated with antibodies to cardiac alpha actinin (green) and co-stained with DAPI (blue; scale bar=10μm). **(D)** Quantification of MYH7-R403Q^+/-^ and control iCM cell area from confocal images stained with alpha actinin. **(E)** Representative immunoblots from MYH7-R403Q^+/-^ and control iCM lysates probed with antibodies to phospho- and total ERK (note: ERK2 is highly phosphorylated in iCM lysates) and phospho- and total AKT as well as GAPDH (loading control). Normalized quantification of **(F)** phospho-AKT to total AKT and **(G)**phospho-ERK2 to total ERK2. Significance was assessed by Student’s t-test (B, D, F, G; *all p<0.05); data are means +/- SEM (error bars) (B, D, F, G).

### RNA sequencing identifies activation of p53 signaling in HCM models

To identify molecular mechanisms of HCM mutations, we applied RNA sequencing to iCM samples from HCM mutations (MYH7-R403Q^+/-^, MYBPC3^+/-^ and MYBPC3-R502W^+/-^) and compared to isogenic controls. We started by analyzing the MYBPC3^+/-^ mutation, tryptophan-792-valine-fs, to determine how MYBPC3 truncation mutations cause HCM. Consistent with nonsense-mediated messenger RNA decay, transcripts from tryptophan-792-valine-fs allele were nearly absent compared to wildtype transcripts (fig.5A). This is in contrast to the missense mutation R502W, which had no evidence of degradation of mutant transcripts. In parallel to changes in MYBPC3 transcript level, total protein content was also reduced in MYBPC3^+/-^ (fig.5B, C) but not R502W^+/-^ (fig.S4A, B) by immunoblotting iCM lysates with an antibody that recognizes the amino-terminus of MYBPC3 proximal to the frameshift mutation. These data support a haploinsufficiency model of HCM-associated MYBPC3 frameshift mutations, and also that R502W is likely a loss-of-function missense mutation.

Next, we analyzed iCM gene transcripts by unsupervised principle component analysis (PCA) to assess sample-to-sample distances. All HCM samples clustered distinctly from isogenic controls, while biological replicates within the same genotype clustered closely (fig.5D, E and table S2). Of note, gene components obtained from PC1 and PC2 include mitochondrial-encoded transcripts (*MT-TP*, *MT-ND6*, *MTATP6P1*, *MT-ND4L*, *MT-ATP8*, *MT-RNR1* and *MT-ND2*), extracellular matrix-related transcripts (*COL3A1*, *FBN1*, *SULF1*, *BGN* and *LGALS1*), sarcomere transcripts (*ACTC1*, *MYL3* and *MYL7)* and p53-related gene targets (*CDKN1A*, *LINC01021*)^31^.

To better define the HCM-associated gene transcript program, we analyzed differentially-expressed transcripts that were common to all HCM iCM models (fig.S4C and table S3). In accord with the close proximity of HCM samples by PCA analysis, differentially-expressed transcripts were also highly shared among HCM models. Of 1050 total upregulated transcripts, 320 were shared by at least two HCM models, and of the 1177 total downregulated transcripts, 419 were shared by at least two HCM models. Of note, the HCM phenotype did not correlate with expression of cardiac chamber-specific markers such as myosin light chain 7 (*MYL7*), myosin light chain 2 (*MYL2*) and potassium channel *HCN4* (fig.S4D). We then used differential expression results to perform pathway analysis in HCM iCM models using Ingenuity Pathway Analysis (IPA). Both p53 and CDKN1A (p21) pathways were predicted to be highly activated in HCM iCMs (fig.5F, table S4) as prioritized by activation Z-score and p-value of overlap. In parallel to transcript levels, p53 and p21 protein levels were increased in HCM iCM lysates (fig.5G-I). In addition, fixed and immunostained HCM CMTs also exhibited increased p53 staining within the nuclei of ACTN2+ iCMs (fig.5J, K). Unexpectedly, these results implicate altered p53 signaling as a common molecular consequence of thick filament HCM mutations.

### HCM iCMs exhibit increased cytotoxicity induced by metabolic stress that can be rescued by p53 inhibition

To understand the role of elevated p53 signaling in HCM iCM models, we first determined whether p53 activation was related to increased iCM stress. In normal growth media, we observed no change in HCM iCM cell death (fig.S4E), which was also consistent with our observation that CMT cell number and tissue cross-sectional area were not changed by HCM mutations. We then considered whether HCM iCMs were more susceptible to stress. Informed by the clinical observation that HCM patients have bio-energetic deficits *in vivo*^32^ as well as the increased expression of mitochondrial gene transcripts in HCM iCMs (fig. 5E), we measured iCM cytotoxicity induced by metabolic stress from glucose removal that has been used by others to induce energy stress^33^. We found that HCM iCM models exhibited elevated cytotoxicity (fig.6A) compared to controls, which was associated with increased ADP:ATP (fig.6B)—a molecular marker of metabolic stress. We next tested whether p53 knockdown by lentiCRISPR could rescue HCM-associated cytotoxicity induced by metabolic stress. P53 knockdown rescued HCM cytotoxicity induced by metabolic stress (fig.6C), and also resulted in reduced CDKN1A transcript levels (fig.6D), which is a transcriptional target of P53.

**Figure 5.**
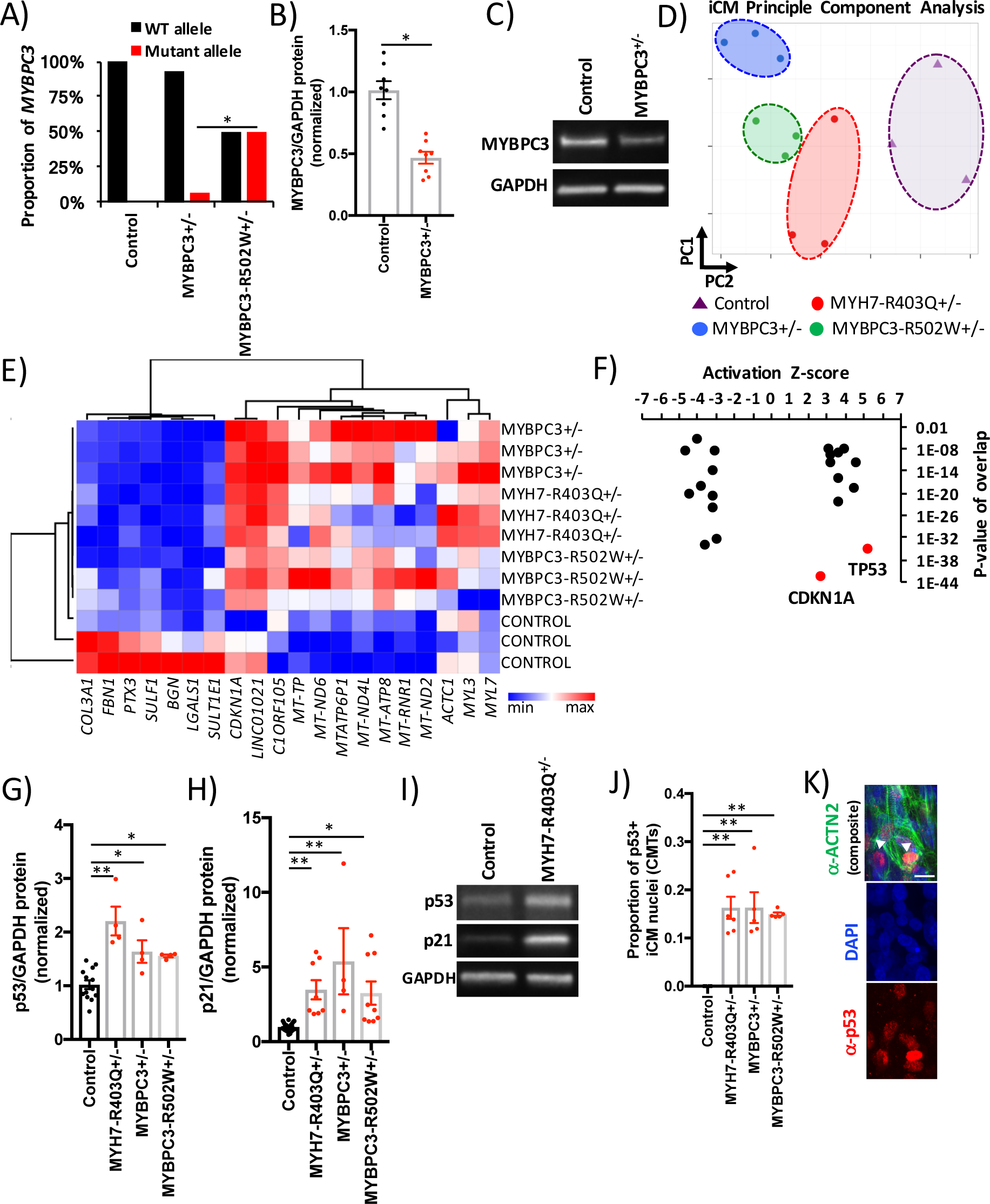
RNA sequencing of HCM and isogenic control iCMs. **(A)** By allele-specific analysis of gene transcripts obtained from isogenic control, MYBPC3^+/-^ and MYBPC3-R502W^+/-^ iCMs; MYBPC3 transcripts were quantified for control (WT; black bar) and mutant (red bar) alleles and shown as proportion of total MYBPC3 expression. (**B**) Densitometry of immunoblots from protein lysates derived from control and MYBPC3^+/-^ iCMs, and probed for MYBPC3 (note: truncated MYBPC3 was not identified) and for protein loading with GAPDH (glyceraldehyde 3-phosphate dehydrogenase; GAPDH). **(C)** Representative immunoblot from (B). (**D**) Principle component analysis (PCA) of RNA transcripts from three biological replicates of isogenic control (purple triangles) and MYBPC3^+/-^ (blue circles), MYBPC3-R502W^+/-^ (green circles) and MYH7-R403Q^+/-^ iCMs (red circles). (**E**) Hierarchical clustering of genes contributing to PC1 and PC2 from (D) and illustrated by heatmap (table S2). (**F**) Differentially-expressed gene transcripts (log_2_FC >0.3 or <-0.3 and FDR<0.1) were analyzed by pathway analysis using Ingenuity Pathway Analysis and identified pathways (black and red dots) were organized by activation z-score and p-value of overlap. (**G**) Densitometry of immunoblots from control and HCM iCM protein lysates, normalized for protein loading (GAPDH) and probed with antibodies to p53 or **(H)** p21 (see representative blots **(I)**). **(J)** Quantification of iCM p53+ nuclei from confocal images of fixed CMTs immunostained with an antibody to p53 (red), cardiomyocyte-specific ACTN2 and DAPI co-stain (see representative image in (**K**) (arrowhead marks p53+ nuclei that co-stain with ACTN2; scale bar=15μm). Significance (*p<0.05 and **p<0.001) was assessed by Fisher’s exact test (A), Student’s t-test (B) or ANOVA (G, H and J); and data are means +/- SEM (error bars) (B, G, H and J).

**Figure 6.**
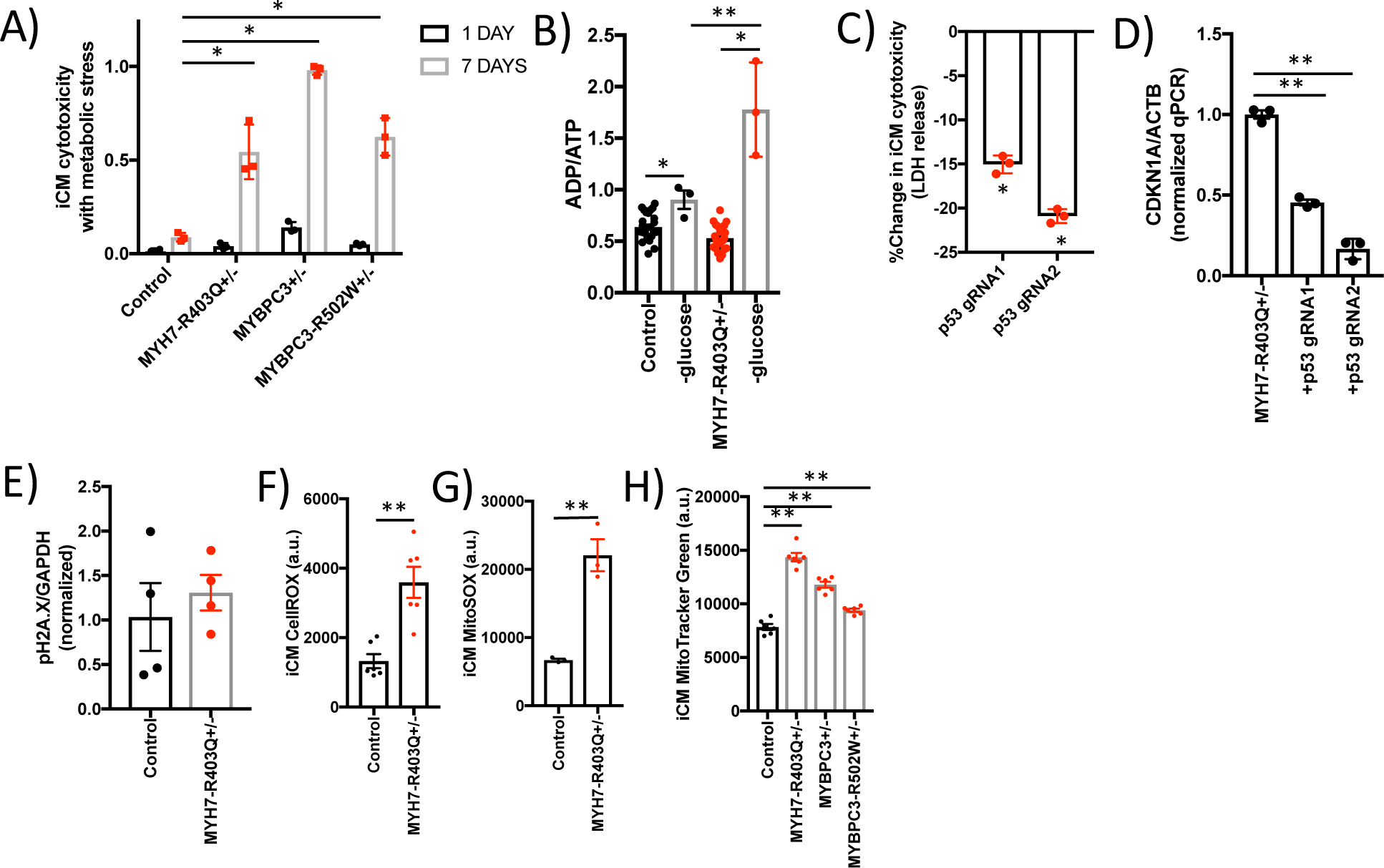
HCM iCMs exhibit a p53-dependent cytotoxicity with metabolic stress that is related to increased oxidative stress. **(A)** Proportion iCM death after one and seven days of metabolic stress induced by growth in glucose-free media. **(B)** ADP:ATP for iCMs cultured in normal growth media and after 24-hours in glucose-free media. **(C)** Change in iCM cytotoxicity measured by LDH release assay upon p53 genetic knockdown using lentiCRISPR encoding two independent gRNAs that target the *TP53* gene. **(D)** Quantitative PCR analysis of p21 (*CDKN1A*) normalized to *ACTB* obtained from cDNA libraries generated from MYH7-R403Q+/- iCMs transduced with lentiCRISPR encoding two independent TP 53 or non-targeted gRNAs. **(E)** Quantification of phosphorylated H2A.X normalized to GAPDH using densitometry analysis of immunoblots from iCM lysates.). **(F)** Quantification of fluorescence (arbitrary units) using FACS analysis of iCMs stained with CellROX green or **(G)** MitoSOX red, a mitochondrial-specific probe for reactive oxygen species. **(H)** Quantification of fluorescence (arbitrary units) using FACS analysis of iCMs stained with MitoTracker Green. Significance (*p<0.05 and **p<0.001) was assessed by ANOVA (A) or Student’s t-test (B-H); and data are means +/- SEM (error bars) (A-H).

To gain insights into linkages between thick filament HCM mutations and p53 activation, we started by measuring known molecular activators of p53 including DNA damage^34^ and oxidative stress^35^. While the levels of phosphorylated histone 2A member X (H2A.X; fig.6E), a marker of DNA damage was not different between R403Q^+/-^ iCMs compared to controls, reactive oxygen species (ROS) levels were increased over three-fold (fig.6F). Because ROS generation is most commonly generated in mitochondria, we also stained iCMs with MitoSOX (fig.6G) and MitoTracker dyes (fig.6H) to quantify mitochondrial-derived ROS and mitochondrial content, respectively. Both of these assays showed increased signal in HCM iCMs compared to controls. Based on these results, we conclude that HCM mutations result in increased oxidative stress that results in a p53-dependent metabolic vulnerability to energy stress.

## Discussion

The genetic basis of HCM, most commonly due to thick filament sarcomere mutations, was identified decades ago, yet the mechanisms that connect sarcomere gene mutations with the HCM phenotype still remain unclear in part because of the lack of a robust human *in vitro* model system to interrogate proximal mechanisms of HCM pathogenesis. Here, we engineered four human HCM models using CRISPR to generate isogenic mutations in two of the most commonly mutated sarcomere genes, and measured the mechanical consequences of these variants in a 3d CMT assay. This approach demonstrated a convergent model of HCM pathogenesis whereby thick filament HCM mutations induce a state of hypercontractility due to both increased twitch force and resting tension. The increased resting tension has particular importance as this component of contraction force is not affected by therapeutics that target only voltage-dependent factors such as the L-type calcium channel.

Our results of hypercontractility are in accord with recent functional studies of HCM-associated thin filament mutant mouse models that support a sarcomere tension model^36^. Thin filament HCM mutations were found to increase tension primarily by increasing myofilament calcium sensitivity as would be expected by changes in troponin complex function, while our findings are consistent with a model whereby thick filament HCM mutations result in hypercontractility independent of changes in calcium handling. Our model is also supported by recent biophysical studies of MHC-β and MYBPC3 interactions that report changes in the accessibility of myosin heads to generate force in the setting of HCM mutations^37^. Our inability to observe previously described alterations in calcium handling may be due to the nature of the variants studied or due to non-cell-intrinsic mechanisms. We propose that calcium handling defects are likely indirect consequences of heart failure, but not directly due to thick filament HCM mutations. In the future, it will be important to test other thick filament variants to identify potential functional heterogeneity, as well as to compare to thin filament CMT phenotypes.

Our study also illustrates how HCM-associated hypercontractility is maladaptive due to increased oxidative stress, which is associated with reduced viability in the setting of metabolic stress. Energy imbalance in HCM patients with significant LVH has been identified in other studies^38^, but our study is the first to implicate an early, cell-intrinsic metabolic vulnerability that results in altered cell survival. While we did not observe an energy imbalance in HCM iCMs in unstressed conditions, this may be secondary to the fetal nature of iCM metabolism that favors glucose rather than fatty acids for energy production.

Finally, we identify that HCM-associated ROS production leads to activation of p53. While p53 has been well studied in the context of cancer and other disorders, it has not been implicated in HCM pathogenesis to our knowledge, and only recently implicated in regulating the cardiac transcriptome^39^. Notably, p53 genetic ablation also protected against heart failure due to pressure-overload, but promoted age-associated cardiac dysfunction^39^. Our results suggest that p53 ablation would be beneficial in HCM, and therefore molecular linkages between sarcomere variants, oxidative stress and p53 activation will need to be addressed in future studies with potential therapeutic implications. Finally, our robust genome-engineered platform using isogenic human iPScs combined with CMT assays demonstrates a robust approach to interrogate HCM pathogenesis, test cardiomyopathy therapeutics, as well as classify sarcomere gene variants of unknown significance.

## Methods

### IPSc culture, iCM differentiation and enrichment

Parental human iPSc line used for all studies was PGP1, a normal control line obtained from Coriell Institute (GM23338), and that has been described previously^18, 20^. This study was approved by the Jackson Laboratory institutional review committee and IRB approval was obtained as part of the Personal Genome Project. All iPSc lines were maintained on Matrigel (Corning)-coated tissue culture plates in mTeSR1 media (Stemcell). IPSc lines were screened for copy number variants using Illumina SNP arrays as previously described^18^. IPScs were differentiated to iCMs by sequential targeting of the WNT pathway as described previously^40^. ICMs were maintained in RPMI with B27 supplement (ThermoFisher) unless otherwise noted. ICMs were enriched by metabolic selection by previously described methods^17^. On day 12 of iCM differentiation, beating iCMs were enriched by adding 4 mM DL-lactate (Sigma) in glucose-free DMEM media (ThermoFisher) for 48 hours. Following selection, enriched iCMs were maintained in RPMI with B27 supplement. Only differentiation batches with >90% troponin T2+ cells by FACS or estimated by morphology were considered to be high purity for further assays. For all assays, iCMs were studied on differentiation day 30-35. For all iCM and CMT assays, at least biological triplicates (differentiation batches) were studied.

### CRISPR/Cas9 iPSc and iCM gene editing

Genetic modifications were generated using scarless iPSc clonal selection by following protocols described previously^18^. For *MYH7* and *MYBPC3* mutation generation, PGP1 iPScs were electroporated with pCAG-eGFP-2A-CAS9 plasmid (obtained from Addgene), a single-stranded mutation-specific 90mer oligonucleotide and an optimized guide RNA plasmid (table S2). After 48 hours, GFP+ iPScs were sorted by flow cytometry (FACS), clonally expanded and Sanger sequenced for genotyping. Guide RNAs were designed and optimized using *in silico* methods that reduce risk of off-target mutations (crispr.mit.edu). For *MYH7* R403Q gene-editing, the corresponding region of *MYH6* was sequenced to verify no off-target *MYH6* mutation. For lentiCRISPR (v2) experiments (Addgene), protocols for gRNA cloning and lentivirus production was obtained from published methods and multiplicity of infection of 3 was used^41^.

### CMT production and force measurements

CMTs were prepared as previously described ^18^. Polydimethylsiloxane (PDMS; Sylgard 184 from Corning) cantilever devices were molded from SU-8 masters, with embedded 1 μM fluorescent microbeads (carboxylate fluospheres; ThermoFisher). PDMS tissue gauge substrates were treated with 0.2% pluronic F127 (Sigma) for 30 minutes to reduce cell-extracellular matrix interactions. ICMs were disassociated using trypsin digestion and mixed with stromal cells (human cardiac fibroblasts; single lot obtained from Lonza), which were pre-treated with 10 μg/ml Mitomycin C (Sigma) to prevent cell proliferation. The number of stromal cells was 7% of the total cell population, which is the quantity necessary for tissue compaction. A suspension of 1.3e^6^ cells within reconstitution mixture containing 2.25 mg/ml collagen I (BD Biosciences) and 0.5 mg/ml human fibrinogen (Sigma) was added to the substrate. We measured CMT function at day seven to allow for tissue compaction and stability of force generation. For quantifying tissue forces, fluorescence images were taken at 25 Hz with an Andor Dragonfly microscope (Andor iXon 888 EMCCD camera with HC PL Fluotar 5X objective mounted on a DMI8 (Leica) microscope that was equipped with a fully-enclosed live-cell environmental chamber (Okolabs)). All tissues were biphasic stimulated at 1 Hz with a C-Pace EP stimulator (IonOptix) and platinum wire electrodes that were separated by 2 cm to the sides of the tissues tested. The displacement of fluorescent microbeads was tracked using the ParticleTracker plug-in in ImageJ (National Institutes of Health). Displacement values were analyzed in Excel (Microsoft) to compute twitch force (dynamic force), resting tension and kinetics. Resting tension was measured by subtracting the resting cantilever position from the cantilever position prior to tissue generation. Cantilever spring constants were computed using the empirically determined elastic modulus of PDMS and the dimensions of the tissue gauge device as described previously^19^. For small molecule treatment, CMTs were treated in Tyrode’s solution for 10 minutes with verapamil (Tocris), blebbistatin (Tocris), pifithrin-*α* (Tocris) or n-acetylcysteine (Sigma) prior to force measurements.

### Calcium (Ca^2+^) imaging using Fluo-4 AM and Indo-1 AM

CMTs were loaded with 4 μM Fluo-4 AM (ThermoFisher) for 60 minutes at 37°C in Tyrode’s solution (140 mM NaCl, 5.4 mM KCL, 1mM MgCl_2_, 10mM glucose, 1.8 mM CaCl_2_ and 10mM HEPES pH 7.4 with NaOH). Following loading, tissues were washed three times in Tyrodes solution. Using a solid-state 50 mW 488 nm laser and GFP filter (emission 525+ 50nm), fluorescence intensities (F/F_o_) were obtained while pacing tissues at 1 Hz. All images were captured at 30 fps on an Andor Dragonfly multimodal microscope. For each tissue, at least four regions of interest (ROIs) were analyzed for changes in Fluo-4 intensity, with the resting fluorescence value F_o_ determined by the average of the first five frames of the video. Background intensity was subtracted from all values. For iCM resting and caffeine-induced calcium quantification, single iCMs were disassociated from monolayer differentiation batches using accutase (ThermoFisher). Disassociated iCMs were loaded with 5 μM Indo-1 AM (ThermoFisher) in Tyrodes solution. ICMs were analyzed using a FACSymphony A5 analyzer (BD Biosciences). After excitation with a 60 mW 355 nm laser, Indo-1 signal ratios were determined using a 450nm dichroic mirror and BUF395 and BUV496 filters. To determine baseline calcium concentration, iCMs were analyzed for 20 seconds and the average Indo-1 ratio was obtained. To determine caffeine-induced calcium levels, a stock of 20 mM caffeine (ThermoFisher) in Tyrodes buffer was subsequently added to reach a 10 mM caffeine concentration, and maximum Indo-1 ratio was determined instantaneously.

### Immunofluorescence, protein quantification and sarcomere analysis

Protein lysates obtained from iCMs were solubilized in RIPA buffer followed by western blotting. For analysis of MYBPC3 protein levels, an antibody that recognizes an n-terminal epitope of MYBPC3 protein was generously provided by Samantha Harris^42^. All other antibodies used for immunoblotting include p53 (DO-7; ThermoFisher), p21 (Cell Signaling), phospho- and total ERK (Cell Signaling), phospho- and total AKT (Cell Signaling), phospho- and total p38 (Cell Signaling), phospho- and total JNK (Cell Signaling), phospho- and total CAMKII (Cell Signaling), pH2A.X (Cell Signaling), GAPDH (Cell Signaling) and TNNI3 (ThermoFisher). For immunofluorescence, iCMs and CMTs were fixed with 4% paraformaldehyde (PFA) for 10 minutes, permeabilized and stained with antibodies to cardiac alpha actinin (Abcam) with DAPI co-stain. All immunofluorescence imaging was completed using a solid-state laser confocal microscope (Andor Dragonfly). CMT sarcomere analysis was done according to established methods^43^. To determine z-disk anisotropy, CMTs were fixed with 4% PFA, permeabilized and immunostained with antibodies directed against Z-disk protein cardiac alpha actinin. Confocal images were obtained and analyzed using a modified ridge detection algorithm in ImageJ (NIH) to produce spatial maps of sarcomere Z-disks and the local orientation with respect to the x-axis of the image. For sarcomere length analysis, CMTs were relaxed in calcium-free buffer, fixed with 4% PFA, permeabilized and immunostained with antibodies directed against Z-disk protein cardiac alpha actinin. Confocal images were analyzed in ImageJ (NIH) using a Z-disk profile plot of greater than three regions of interest per CMT. ICM cell area was measured in iCMs that were cultured on glass coverslips, fixed with 4% PFA, permeabilized, and similarly immunostained. ICM cell area were measured using ImageJ.

### Cell viability and oxidative stress assays

For metabolic stress assays, iCMs were cultured at sub-confluence in 6-well plates coated with fibronectin (10ug/ml; Corning) at a density of 100,000 cells/well. Prior to metabolic stress conditions, cells were maintained in RPMI with B27 supplement. To induce nutrient stress, cells were maintained in glucose-free DMEM media (ThermoFisher) with B27 supplement for up to 7 days. To quantify iCM death, either LDH released from dead cells was analyzed from conditioned media using an LDH release assay (ThermoFisher), or viable cells were counted using live-cell imaging and a viability stain (MitoTracker Red). ICM conditioned media was analyzed in a 96-well plate with a Synergy plate reader (BioTek) to quantify absorbance at 490 nm and 680 nm. For ADP/ATP ratios, 20,000 iCMs were plated on fibronectin-coated (10ug/ml; Corning) 96-well plates and analyzed by a bioluminescent ADP/ATP assay (Biovision).

### RNA sequencing, quantitative PCR and computational analyses

For iCM RNA sequencing experiments, total RNA was isolated from day 30-35 iCMs using Trizol (ThermoFisher). CDNA was constructed using Superscript III First-Strand synthesis (ThermoFisher). For each sample, total RNA from biological triplicates by differentiation batch was collected and sequenced. RNA sequencing libraries were generated using the TruSeq Stranded mRNA library preparation kit (Illumina). RNA sequencing libraries were sequenced on a HiSeq 2500 v4 SBS platform 2×100bp reads (Illumina). Sequences were aligned with STAR^44^ to the hg38 human genome. For differential expression, DESeq2 (Bioconductor) was used. Heatmap analysis and hierarchical clustering were performed using the Morpheus tool (Broad Institute). For quantitative PCR analysis, iCM total RNA was isolated and processed as above. Gene-specific PCR primers were identified from the literature or designed using Primer3 (see table S2 for sequences) and transcripts were quantified using Fast SYBR Green (Applied Biosystems) on an ViiA7 Real-Time PCR system (Applied Biosystems).

### AFM Modulus Quantification

To measure the elastic modulus of the PDMS used for tissue gauge production, indentation measurements were performed using Atomic Force Microscopy (AFM) (Asylum Research MFP-3D, Santa Barbara, Ca). Using a colloidal silica spherical probe (AppNano FORT SiO_2_-A), a 16×16 array of force-distance curves was acquired evenly spaced across a 2×2µm area for each measurement. Before and after these force-volume experiments, identical force-distance measurements on an incompressible surface were performed with a steel disk to calibrate the system sensitivity and to determine the spring constant for the force-transducing cantilever (1.75 nN/nm of deflection, within the range of specifications for these commercial probes). All 256 force-distance curves for each location studied were analyzed based on the Johnson-Kendall-Roberts model of elastic contact. Such JKR mechanics accommodate large adhesion energies between an indenting probe and sample^45, 46^, both expected and observed for these studies with PDMS. This incorporates the following experimental parameters: the indenter possesses a spherical tip geometry with a radius of 2.5 µm; the probe modulus appropriate for the colloidal silica sphere is 68.0 GPa; and this probe exhibits a Poisson ratio of 0.19. A Poisson ratio of .33 is conventionally assumed for the specimen. For each of the samples, force-volume maps were completed at three representative locations. The average elastic modulus from the resulting 768 individual indentations per specimen is reported.

## Acknowledgements

We thank Anthony Carcio for contributions to the flow cytometry experiments, and Qianru Yu for expertise in confocal microscopy. We also thank Bo Reese for RNA sequencing technical expertise. We thank Samantha Harris for her generous contribution of MYBPC3 antibody.

## Sources of Funding

Funding for this work was supported by the National Heart, Lung, and Blood Institute at the NIH (HL125807 and HL142787).

## Disclosures

J.T.H. received a research grant from Myokardia, Inc. to study dilated cardiomyopathy CMT models. Myokardia is developing small molecule direct myosin activators/inhibitors for cardiomyopathy.

## Author Contributions

R.C., K.T., A.L., E.M., Y.C., and J.T.H. designed and performed iPSC and iCM research, analyzed and interpreted data. J.T.H. wrote the manuscript with assistance from R.C. and K.T.. R.C. designed and engineered cardiac tissue devices. A.L., F.L., A.P., K.T. and R.C. generated cells for this study. K.A. and B.D.H. performed AFM analysis of PDMS devices.

